# Structural basis for potent neutralization of SARS-CoV-2 and role of antibody affinity maturation

**DOI:** 10.1101/2020.06.12.148692

**Authors:** Nicholas K. Hurlburt, Yu-Hsin Wan, Andrew B. Stuart, Junli Feng, Andrew T. McGuire, Leonidas Stamatatos, Marie Pancera

## Abstract

SARS-CoV-2 is a betacoronavirus virus responsible for the COVID-19 pandemic. Here, we determined the X-ray crystal structure of a potent neutralizing monoclonal antibody, CV30, isolated from a patient infected with SARS-CoV-2, in complex with the receptor binding domain (RBD). The structure reveals CV30’s epitope overlaps with the human ACE2 receptor binding site thus providing the structural basis for its neutralization by preventing ACE2 binding.

## Main

COVID-19 was declared a pandemic in March 2020 by the World Health Organization^1^. As of June 11^th,^ 2020, there were ∼ 7.4 M infections and over 415,000 deaths worldwide^2^. It is caused by a coronavirus of the beta family, named SARS-CoV-2^3^, as it is closely related to SARS-CoV^4^. Their genomes share 80% identity and they utilize angiotensin-converting enzyme 2 (ACE2) as receptor for entry^5-11^. Viral entry depends on the SARS-CoV-2 spike glycoprotein, a class I fusion protein comprised of two subunits, S1 and S2. S1 mediates ACE2 binding through the receptor binding domain (RBD), while the S2 subunit mediates fusion. Overall the spike shares 76% amino acid sequence homology with SARS^4^. High resolutions structures of the SARS-CoV-2 stabilized spike in the prefusion revealed that the RBD can be seen in a ‘up’ or ‘down’ conformation^5,6^.It’s been shown that some of the neutralizing antibodies bind the RBD in the ‘up’ conformation similar to when the ACE2 receptor binds^12^. Currently there is no vaccine available to prevent SARS-CoV-2 infection and highly effective therapeutics have not been developed yet either. The host immune response to this new coronavirus is also not well understood. We, and others, sought to characterize the humoral immune response from infected COVID-19 patients^12-14^. Recently, we isolated a neutralizing antibody, named CV30, which binds the receptor binding domain (RBD), neutralizes with 0.03 μg/ml and competes binding with ACE2^15^. However, the molecular mechanism by which CV30 blocked ACE2 binding was unknown. Herein, we present the 2.75 Å crystal structure of SARS-CoV-2 RBD in complex with the Fab of CV30 (Extended Data Table 1).

CV30 binds almost exclusively to the concave ACE2 binding epitope (also known as the receptor binding motif (RBM)) of the RBD using all six CDR loops with a total buried surface area of ∼1004 Å^2^, ∼750 Å^2^ from the heavy chain and ∼254 Å^2^ from the kappa chain (Fig. 1A). 20 residues from heavy chains and 10 residues from the kappa chain interact with the RBD, forming 13 and 2 hydrogen bonds, respectively (Fig. 1C and Extended Data Table 2). There are 29 residues from the RBD that interact with CV30, 19 residues with the heavy chain, 7 residues with the light chain, and 3 residues with both (Extended Data Table 2). Of the 29 interacting residues from the SARS-CoV-2 RBD, only 16 are conserved in the SARS-CoV S protein RBD (Fig. 2c), which could explain the lack of cross-reactivity of CV30 to SARS-CoV S^15^. The CV30 heavy chain is minimally mutated with only a two-residue change from the germline and both of these residues (Val27-Ile28) are located in the CDRH1 and form nonpolar interactions with the RBD. We reverted these residues to germline to assess their role. Interestingly, the germline CV30 (glCV30) antibody bound to RBD with ∼100-fold lower affinity (407 nM affinity) (Fig 1d and Extended Data Table 3) compared to CV30 (3.6 nM^15^) with a very large difference in the off-rate. glCV30 neutralized SARS-CoV-2 with ∼500-fold difference with an IC50 of 16.5 vs 0.03 μg/mL for CV30 (Fig. 1e). Val27 forms a weak non-polar interaction with the RBD Asn487 and sits in a pocket formed by CDRH1 and 3. Although it is unclear, Phe27 presents in glCV30 could change the electrostatic environment. The Ile28 sidechain forms non-polar interactions with the RBD Gly476-Ser447, particularly the Cγ atom, which the glCV30 Thr would be incapable of making. Thus, minimal affinity maturation of CV30 significantly impacted the ability of this mAb to neutralize SARS-CoV-2.

**Figure 1.**
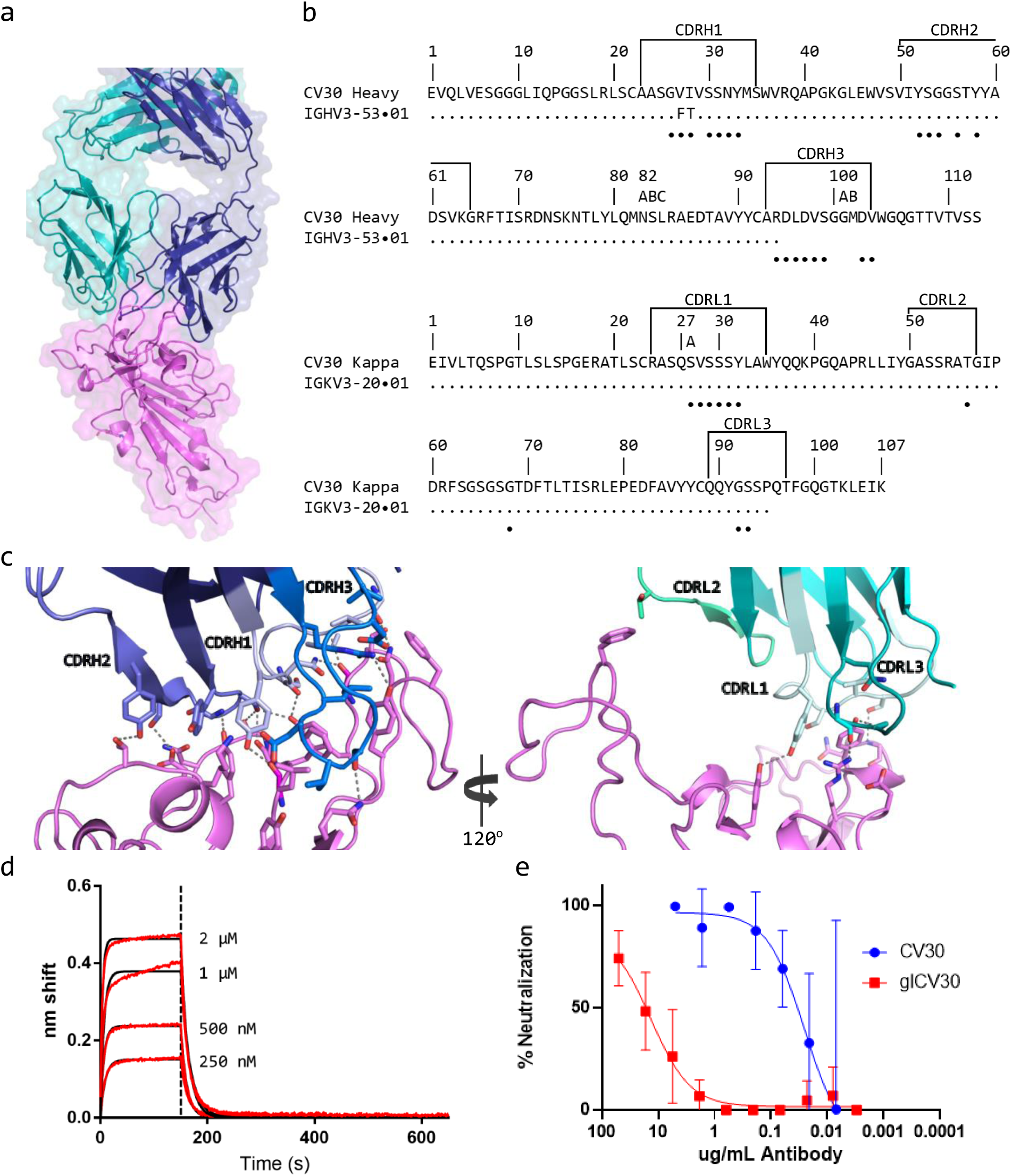
Overall structure of CV30 Fab in complex with SARS-CoV-2 RBD and kinetics of glCV30. **a**. Structure is shown in cartoon with surface representation shown in transparency. CV30 heavy chain is shown in dark blue and light chain in light blue. RBD is shown in pink. **b**. Sequence alignment of CV30 heavy and light chains with germline genes. Black circles underneath the sequence indicate residues that interact with the RBD. **c**. Details of the interactions of the heavy (left) and light (right) chains with the RBD. CDRs are labeled and colored as shown. Residues that interacts are shown as sticks and Hydrogen bonds are shown in dotted lines. **d**. Kinetics of glCV30 binding to RBD measured by BLI. **e**. glCV30 and CV30 neutralization of SARS-CoV-2 pseudovirus.

**Figure 2.**
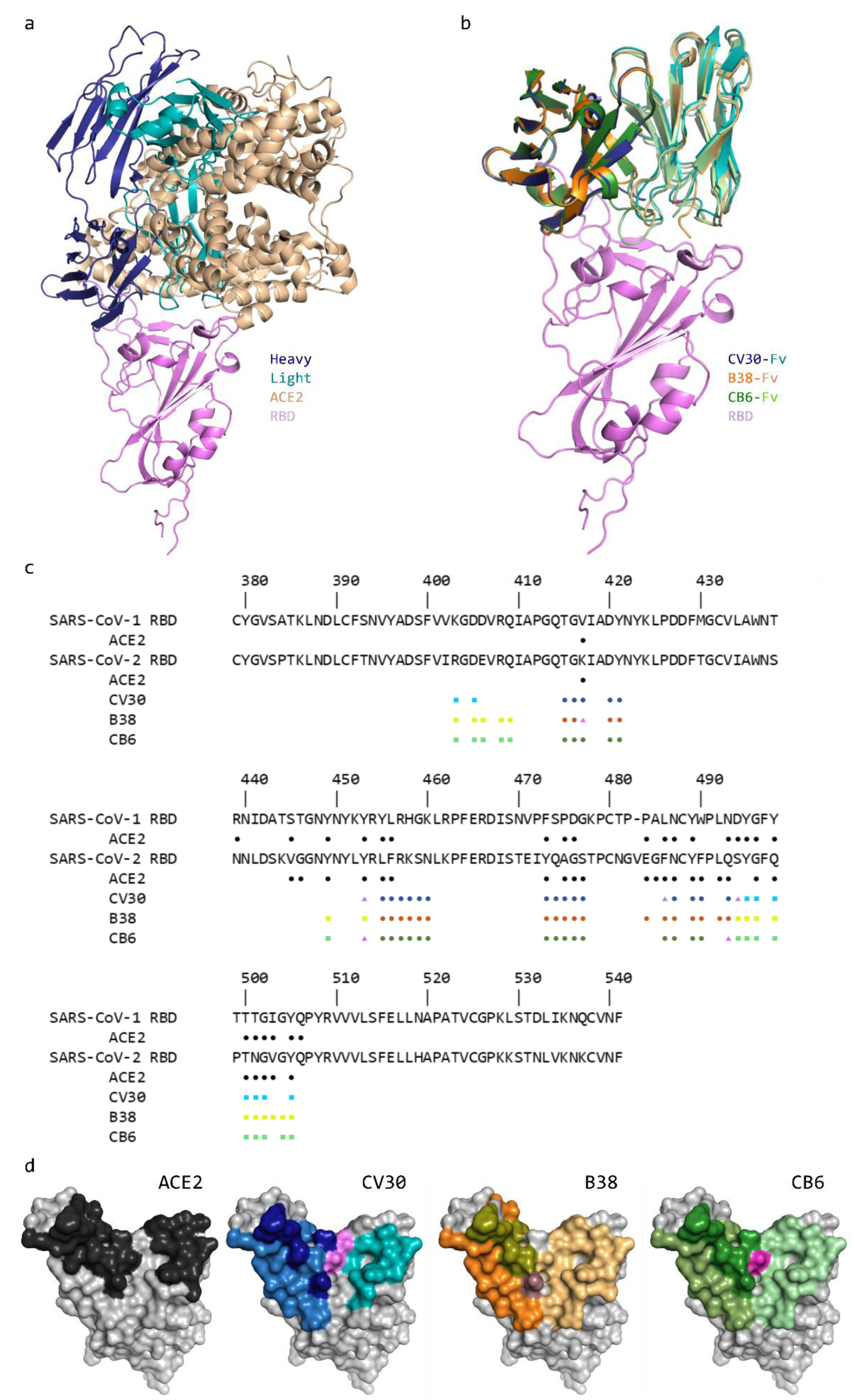
Comparison of the CV30 epitope against ACE2 and other neutralizing antibodies. **a**. Structural overlay of ACE2/RBD complex with CV30/RBD complex. **b**. Structural alignment of the variable domains of CV30, B38, and CB6. **c**. Sequence alignment of SARS-CoV RBD and SARS-CoV-2 RBD. The residues that interact with ACE2 are indicated by the black circles. Residues that interact with CV30, B38, and CB6 are indicated by the colored squares (light chain interactions), circles (heavy chain interactions), or triangles (interactions with both chains). **d**. Surface representation of the RBD with the binding epitope colored. Light chain interactions are the lightest color, heavy chain interactions are next lightest, and CDRH3 specific interactions are darkest, and interacting with both heavy and light chain is purple.

CV30 competes with ACE2 for binding to the RBD^15^ and we therefore examined the structural mechanism of the receptor blocking by superimposing the SARS-CoV-2 RBD/ACE2 complex (PDB: 6LZG)^9^ with the CV30 Fab/RBD complex. The structure of the RBD was used to align the two complexes and showed that CV30 binding did not induce any conformational changes in the RBD from the ACE2-bound complex. The aligned RBD had a RMSD of 0.353 Å over 166 C_α_ atoms. The structure reveals that the CV30 epitope overlaps almost completely with the ACE2 epitope. A total of 26 residues of the SARS-CoV-2 RBD interact with hACE2, CV30 binds to 19 of these residues (Fig. 2A), indicating that CV30 neutralizes the virus by preventing the binding of ACE2 to RBD by direct steric interactions.

Recently, the structure of two potent neutralizing anti-RBD antibodies were published, B38 and CB6^12,14^. CV30 shares a similar germline heavy chain V-genes but all three have diverse germline kappa V-genes (CV30 is IGKV3-20*01, B38 is IGKV1-9*01, CB6 is IGKV1-39*01, Extended Data Fig. 1). Both CV30 and B38 use IGHV3-53*01 while CB6 uses IGHV3-66*01, which is only one amino acid different than 3-53*01 (Val12 which does not make contact with the epitope). CV30 and CB6 each have higher affinities, 3.6 nM and 2.5 nM, respectively, than B38, 70.1 nM^12,14,15^. Differences in affinity translate into differences in neutralization potency (the IC50s for CV30 and CB6 are 0.03 and 0.036 μg/mL, respectively, and that of B38 is 0.177 μg/mL). Interestingly, Thr28 was also mutated from germline to Ile in B38 but Phe27 was not. CB6 lacks both mutations found in CV30. Differences in other regions of the antibody, such as the CDRH3 and light chain are likely responsible for the overall potency all these antibodies (see below). To investigate the binding mechanism of the three antibodies, a superposition of the structures was created. All three bind in a nearly identical manner with the same angle of approach and similar footprints (Fig. 2b). The alignment of the Fv regions of B38 and CB6 to the Fv region of CV30 had a RMSD of 0.240Å over 100 C_α_ atoms and 0.329Å over 98 C_α_ atoms, respectively. Mapping the binding interactions of the RBD to each of the antibodies reveals a close overlap in the binding mechanism (Fig. 2c-d). The footprint of the heavy chain is nearly identical, as expected from the shared germline V-gene and sequence similarity. CV30 and CB6 both have longer CDRH3 and bind with higher buried surface area, ∼263 and ∼251 Å^2^, respectively, than B38 (∼203 Å^2^) (Fig. 2d, Extended Data Fig. 1). The large difference is in the light chain. CV30 has the smallest binding interaction at ∼254 Å^2^, B38 has the largest interaction at ∼497 Å^2^ and then CB6 at ∼354Å^2^. One of the more interesting findings was the interaction of Thr56 in the CV30 CDRK2 which reaches across the RBD and interacts Phe486, an interaction that is not found in the other two antibodies (Extended Data Fig. 1).

In conclusion, our structure indicates that potent neutralizing antibodies against SARS-CoV-2 bind the receptor binding motif in the RBD, overlapping the ACE2 binding site, but recognize residues that are specific for SARS-CoV-2 only, thus explaining the lack of cross neutralization with SARS-CoV. It is noteworthy that potently neutralizing antibodies isolated from multiple individuals use the same or similar VH gene to target their epitope. Additionally, the minimal affinity maturation observed 21 days after infection in the VH gene of CV30 showed ∼100-500-fold increase in affinity and neutralization potency, indicating that further affinity maturation may increase potency and potential cross-reactivity. Our studies indicate that the RBD is a promising target for vaccine design and that these potently neutralizing antibodies should be explored as a treatment for COVID-19 infection.

## Methods

### Recombinant Protein Expression and Purification

The plasmid encoding the receptor binding domain of SARS-CoV-2 spike protein fused to a monomeric Fc (pαH-RBD-Fc) has been previously described^5^ and was a gift from Dr. Jason McLellan.

1L of 293SGlycoDelete cells^16^ were cultured to a density of 1 million cells/mL and transiently transfected with 500μg of pαH-RBD-Fc using 2 mg of polyethylenimine (PEI, Polysciences, Cat# 24765). Cultural supernatant was harvested 6 days post-transfection by centrifugation and sterile filtered using a 0.22μm vacuum filter. The RBD was purified using protein A agarose resin (GoldBio, Cat# P-400) and cleaving the Fc domain using HRV3C protease (made in house) on-column. The eluate containing the RBD was further purified by SEC using a HiLoad 16/600 Superdex 200 pg column (GE Healthcare) column pre-equilibrated in 2mM Tris-HCl, pH 8.0, 200mM NaCl. Protein was aliquoted, flash frozen, and stored at -80°C until needed.

500mL of 293EBNA cells were cultured to a density of 1 million cells/mL and transiently transfected with 125μg each of CV30 Heavy and Kappa chains using 1 mg of PEI. Cultural supernatant was harvested 6 days post-transfection by centrifugation and sterile filtered using a 0.22μm vacuum filter. IgG was purified using protein A agarose resin and eluted using Pierce IgG Elution Buffer (Thermo Scientific, Cat# 21004). Eluate was pH adjusted to 7.5 using 1M HEPES, pH 7.5. IgG was further purified by SEC using a HiLoad 16/600 Superdex 200 pg column. Antigen binding fragment (Fab) was generated by incubating IgG with LysC (New England Biolabs, Cat# P8109S) at a ratio of 1μg LysC per 10mg IgG at 37°C for 18hrs. Fab unexpectedly stuck to protein A resin and was eluted as mixture of Fab, undigested IgG, and digested Fc product using the IgG elution buffer. Fab and Fc product was purified by SEC. The CV30-Fab and SARS-CoV-2 RBD complex was obtained my mixing Fab and Fc product with a 2-fold molar excess of RBD and incubated for 90min at RT with nutation followed by SEC. The complex was verified by SDS-PAGE analysis.

### Crystal Screening and Structure Determination

The complex was concentrated to 10mg/mL for initial crystal screening by sitting-drop vapor-diffusion in the MCSG Suite (Anatrace) using a NT8 drop setter (Formulatrix). Diffracting crystals were obtained in a mother liquor (ML) containing 0.2M (NH4) Citrate, tribasic, pH 7.0 and 12% (w/v) PEG 3350. The crystals were cryoprotected by soaking in ML supplemented with 30% (v/v) ethylene glycol. Diffraction data was collected at Advanced Photon Source (APS) SBC 19-ID at a 12.662 keV. The data set was processed using XDS^17^ to a resolution of 2.75Å. The structure of the complex was solved by molecular replacement using Phaser^18^ with a search model of SARS-CoV-2 RBD (PDBid: 6lzg)^9^ and the Fab structure (PDBid: 5i1e)^19^ divided into Fv and Fc portions. Remaining model building was completed using COOT^20^ and refinement was performed in Phenix^21^. The data collection and refinement statistics are summarized in Extended Data Table 1. Structural figures were made in Pymol.

### BLI

For kinetic analyses glCV30 was captured on anti-Human IgG Fc capture (AHC) sensors at a concentration of 20 μg/mL and loaded for 100s. After loading, the baseline signal was then recorded for 1min in KB. The sensors were immersed into wells containing serial dilutions of purified SARS-CoV-2 RBD in KB for 150s (association phase), followed by immersion in KB for an additional 600s (dissociation phase). The background signal from each analyte-containing well was measured using VRC01 IgG control reference sensors and subtracted from the signal obtained with each corresponding glCV30 loaded sensor. Kinetic analyses were performed at least twice with an independently prepared analyte dilution series. Curve fitting was performed using a 1:1 binding model and the ForteBio data analysis software. Mean kon, koff values were determined by averaging all binding curves that matched the theoretical fit with an R2 value of ≥0.98.

### Neutralization Assay

HIV-1 derived viral particles were pseudotyped with full length wildtype SARS-CoV-2 S^22^. Briefly, plasmids expressing the HIV-1 Gag and pol (pHDM540 Hgpm2), HIV-1Rev (pRC-CMV-rev1b), HIV-1 Tat (pHDM-tat1b), the SARS-CoV-2 spike (pHDM-SARS-CoV-2 Spike) and a luciferase/GFP reporter (pHAGE-CMV-Luc2-IRES542 ZsGreen-W) were co-transfected into 293T cells at a 1:1:1:1.6:4.6 ratio using 293 Free transfection reagent according to the manufacturer’s instructions. 72 hours later the culture supernatant was harvested, clarified by centrifugation and frozen at -80°C.

293 cells stably expressing ACE2 (HEK-293T-hACE2) were seeded at a density of 4×10^3^ cells/well in a 100 µL volume in 96 well flat bottom tissue culture plates. The next day, CV30 and germline CV30 were serially diluted in 30 µL of cDMEM in 96 well round bottom 27 plates in triplicate. An equal volume of viral supernatant diluted to result in 2×10^5^ luciferase units was added to each well and incubated for 60 min at 37 °C. Meanwhile 50 μL of cDMEM containing 6 µg/mL polybrene was added to each well of 293T-ACE2 cells (2 µg/mL final concentration) and incubated for 30 min. The media was aspirated from 293T-ACE2 cells and 100 µL of the virus-antibody mixture was added. The plates were incubated at 37°C for 72 hours. The supernatant was aspirated and replaced with 100 μL of Steadyglo luciferase reagent (Promega). 75 µL was then transferred to an opaque, white bottom plate and read on a Fluorskan Ascent Fluorimeter. Control wells containing virus but no antibody (cells + virus) and no virus or antibody (cells only) were included on each plate.

% neutralization for each well was calculated as the RLU of the average of the cells + virus wells, minus test wells (cells +mAb + virus), and dividing this result difference by the average RLU between virus control (cells+ virus) and average RLU between wells containing cells alone, multiplied by 100. The antibody concentration that neutralized 50% of infectivity (IC50) was interpolated from the neutralization curves determined using the log(inhibitor) vs. response -- Variable slope (four parameters) fit using automatic outlier detection in Graphpad Prism Software.

## Data availability

Coordinates and structure factors for CV30 Fab-SARS-CoV-2 RBD complex have been deposited in the Protein Data Bank (PDB) under the accession code 6XE1.

## Acknowledgments

This work was supported by generous donations to Fred Hutch COVID-19 Research Fund. We thank Dr. McLellan for providing the SARS-CoV-2 RBD plasmid. We thank the J. B. Pendleton Charitable Trust for its generous support of Formulatrix robotic instruments. Results shown in this report are derived from work performed at Argonne National Laboratory, Structural Biology Center (SBC), ID-19, at the Advanced Photon Source. SBC-CAT is operated by UChicago Argonne, LLC, for the U.S. Department of Energy, Office of Biological and Environmental Research under contract DE-AC02-06CH11357.

## Author contribution

N.K.H, A.T.M., L.S and M.P. conceived the project. N.K.H, A.T.M., L.S and M.P designed the experiments. J.F. cloned the plasmids. N.K.H and A.B.S. expressed and purified the proteins. N.K.H. crystallized proteins, collected and processed the diffraction data, and solved the crystal structure. N.K.H and A.J.M. performed kinetic experiments. Y-H. W. and A.J.M performed neutralization assay. N.K.H, A.T.M., L.S and M.P. analyzed and discussed data. N.K.H and M.P. wrote the original manuscript draft. N.K.H, A.T.M., L.S and M.P. reviewed and edited the manuscript.

