## Supplemental Data for "Structural basis for potent neutralization of SARS-CoV-2 and role of antibody affinity maturation"

**Supplemental Table 1**. Data collection and refinement statistics for crystal structure

|  | **CV30 Fab with SARS-CoV-2 RBD** |
| --- | --- |
| **Data collection** |  |
| Space group | P6_3_ |
| Cell dimensions |  |
| *a*, *b*, *c* (Å) | 147.44, 147.44, 89.38 |
| *α, β, γ* (°) | 90, 90, 120 |
| Resolution (Å) | 48.26 – 2.75 (2.85 – 2.75) |
| *R*_merge_^a^ | 0.031 (0.4917) |
| <I/σ(I)> | 13.01 (1.45) |
| CC_1/2_ | 0.999 (0.573) |
| Completeness | 99.93 (99.97) |
| Redundancy | 2.0 (2.0) |
| **Refinement** |  |
| Resolution (Å) | 48.26 – 2.75 (2.85 – 2.75) |
| No. unique reflections | 28886 (2862) |
| *R*_work_^b^/*R*_free_^c^ | 21.14/23.92 (32.83/35.35) |
| No. atoms | 4917 |
| Protein | 4880 |
| Water | 23 |
| Ligand | 14 |
| B-factors (Å^2^) | 78.69 |
| Protein | 78.70 |
| Water | 64.57 |
| Ligand | 98.31 |
| RMS bond length (Å) | 0.003 |
| RMS bond angle (°) | 0.63 |
| **Ramachadran Plot Statistics^d^** | |
| Residues | 638 |
| Most Favored region | 95.08 |
| Allowed Region | 4.60 |
| Disallowed Region | 0.32 |
| Clashscore | **3.21** |
| **PDB ID** | **6XE1** |

^a^ R_merge_ = [∑_h_∑_i_|*I*_h_ – *I*_hi_|/∑_h_∑_i_*I*_hi_] where *I*_h_ is the mean of *I*_hi_ observations of reflection *h*. Numbers in parenthesis represent highest resolution shell. ^b^ R_factor_ and ^c^ R­_free_ = ∑||F_obs_| - |F_calc_|| / ∑|F_obs_| x 100 for 95% of recorded data (R_factor_) or 5% data (R_free_). ^d^ MolProbity reference

**Supplemental Table 2.** Summary of interactions between SARS-CoV-2 RBD and CV30 Fab (from PISA web server, [www.ebi.ac.uk](http://www.ebi.ac.uk))

a. Detailed interactions of SARS-CoV-2 RBD and CV30 Fab Heavy Chain

| **SARS-CoV-2 RBD** | **HSDC** | **ASA** | **BSA** |  | **CV30 Heavy** | **HSDC** | **ASA** | **BSA** |
| --- | --- | --- | --- | --- | --- | --- | --- | --- |
| E:THR 415 | H | 100.21 | 36.26  \|\|\|\| |  | H:GLY  26 |  | 54.70 | 31.89  \|\|\|\|\|\| |
| E:GLY 416 |  | 21.15 | 16.96  \|\|\|\|\|\|\|\|\| |  | H:VAL  27 |  | 10.36 | 8.54  \|\|\|\|\|\|\|\|\| |
| E:LYS 417 | H | 111.60 | 64.06  \|\|\|\|\|\| |  | H:ILE  28 | H | 91.56 | 48.86  \|\|\|\|\|\| |
| E:ASP 420 | H | 21.38 | 19.15  \|\|\|\|\|\|\|\|\| |  | H:SER  30 |  | 26.64 | 2.94  \|\| |
| E:TYR 421 | H | 52.60 | 50.71  \|\|\|\|\|\|\|\|\|\| |  | H:SER  31 | H | 71.94 | 61.25  \|\|\|\|\|\|\|\|\| |
| E:TYR 453 |  | 40.70 | 20.50  \|\|\|\|\|\| |  | H:ASN  32 | H | 23.41 | 21.21  \|\|\|\|\|\|\|\|\|\| |
| E:LEU 455 | H | 46.75 | 46.75  \|\|\|\|\|\|\|\|\|\| |  | H:TYR  33 | H | 92.61 | 83.33  \|\|\|\|\|\|\|\|\| |
| E:PHE 456 |  | 64.15 | 57.70  \|\|\|\|\|\|\|\|\| |  | H:TYR  52 | H | 79.87 | 62.13  \|\|\|\|\|\|\|\| |
| E:ARG 457 |  | 49.01 | 6.19  \|\| |  | H:SER  53 | H | 48.87 | 40.00  \|\|\|\|\|\|\|\|\| |
| E:LYS 458 | H | 146.36 | 42.74  \|\|\| |  | H:GLY  54 |  | 81.91 | 57.87  \|\|\|\|\|\|\|\| |
| E:SER 459 |  | 67.54 | 6.36  \| | | H:SER  56 | H | 53.50 | 42.56  \|\|\|\|\|\|\|\| |
| E:ASN 460 |  | 93.25 | 30.92  \|\|\|\| |  | H:TYR  58 | H | 107.90 | 42.35  \|\|\|\| |
| E:TYR 473 | H | 45.03 | 29.11  \|\|\|\|\|\|\| |  | H:ARG  94 | H | 46.71 | 46.71  \|\|\|\|\|\|\|\|\|\|\| |
| E:GLN 474 |  | 63.55 | 3.44  \| | | H:ASP  95 |  | 34.02 | 1.23  \| |
| E:ALA 475 | H | 56.82 | 56.33  \|\|\|\|\|\|\|\|\|\| |  | H:LEU  96 |  | 53.68 | 47.15  \|\|\|\|\|\|\|\|\| |
| E:GLY 476 |  | 20.93 | 16.74  \|\|\|\|\|\|\|\| |  | H:ASP  97 |  | 93.71 | 42.36  \|\|\|\|\| |
| E:SER 477 |  | 110.57 | 29.70  \|\|\| |  | H:VAL  98 |  | 146.30 | 50.39  \|\|\|\| |
| E:PHE 486 |  | 174.38 | 61.15  \|\|\|\| |  | H:SER  99 | H | 94.68 | 60.64  \|\|\|\|\|\|\| |
| E:ASN 487 | H | 45.99 | 32.88  \|\|\|\|\|\|\|\| |  | H:ASP 101 |  | 72.77 | 9.56  \|\| |
| E:TYR 489 | H | 81.63 | 54.85  \|\|\|\|\|\|\| |  | H:VAL 102 |  | 22.10 | 4.85  \|\|\| |
| E:GLN 493 | H | 75.64 | 29.87  \|\|\|\| |  |  |  |  |  |
| E:SER 494 |  | 44.70 | 0.25  \| |  |  |  |  |  |

b. Detailed interaction of SARS-CoV-2 RBD and CV30 Fab Light chain

| **SARS-CoV-2 RBD** | **HSDC** | **ASA** | **BSA** |  | **CV30 Light** | **HSDC** | **ASA** | **BSA** |
| --- | --- | --- | --- | --- | --- | --- | --- | --- |
| E:TYR 453 | H | 40.70 | 20.20  \|\|\|\|\| | | L:SER  27A |  | 88.22 | 52.82  \|\|\|\|\|\| |
| E:PHE 486 |  | 174.38 | 7.35  \| | | L:VAL  28 |  | 8.77 | 7.19  \|\|\|\|\|\|\|\|\| |
| E:SER 494 |  | 44.70 | 0.49  \| | | L:SER  29 |  | 67.12 | 52.97  \|\|\|\|\|\|\|\| |
| E:TYR 495 |  | 9.83 | 4.43  \|\|\|\|\| | | L:SER  30 |  | 45.62 | 2.21  \| |
| E:GLY 496 |  | 39.99 | 17.46  \|\|\|\|\| | | L:SER  31 |  | 67.76 | 6.62  \| |
| E:GLN 498 |  | 68.21 | 2.70  \| | | L:TYR  32 | H | 95.94 | 58.65  \|\|\|\|\|\|\| |
| E:THR 500 |  | 125.50 | 7.00  \| | | L:THR  56 |  | 142.51 | 7.77  \| |
| E:ASN 501 |  | 38.08 | 15.54  \|\|\|\|\| | | L:GLY  68 |  | 27.96 | 1.17  \| |
| E:GLY 502 |  | 39.16 | 25.23  \|\|\|\|\|\|\| | | L:GLY  92 |  | 42.55 | 32.49  \|\|\|\|\|\|\|\| |
| E:TYR 505 | H | 124.11 | 110.81  \|\|\|\|\|\|\|\|\| |  | L:SER  93 | H | 47.50 | 24.38  \|\|\|\|\|\| |

**HSDC H**ydrogen/**D**isulfide bond, **S**alt bridge or **C**ovalent link; **ASA**  Accessible Surface Area, Å²;

**BSA**  Buried Surface Area, Å²  ||||   Buried area percentage, one bar per 10%

**Supplemental Table 3.** Kinetic analysis of glCV30 binding to SARS-CoV-2 RBD

| Ligand | Analyte | K_D_ (M X 10^-9^) | k_on_ (1/Ms) x10^5^ | k_on_ error x10^4^ | k_off_ (1/s) x10^-2^ | K_off_ error x10^-3^ |
| --- | --- | --- | --- | --- | --- | --- |
| glCV30 IgG | SARS-CoV-2 RBD | 407 | 1.83 | 1.40 | 7.46 | 1.12 |

**Supplemental Figure 1.** Sequence alignment of CV30, B38, and CB6 heavy and light chain variable regions. Residues interacting with RBD are indicated with ▪. Conserved residues are indicated with * and similar residues are shown with : or . .

```
CV30_Heavy      EVQLVESGGGLIQPGGSLRLSCAASGVIVSSNYMSWVRQAPGKGLEWVSVIYSGGSTY	58
                                         ▪▪▪ ▪▪▪▪                  ▪▪▪ ▪ ▪
B38_Heavy       EVQLVESGGGLVQPGGSLRLSCAASGFIVSSNYMSWVRQAPGKGLEWVSVIYSGGSTY	58
                                         ▪▪▪ ▪▪▪▪                  ▪▪▪ ▪▪▪
CB6_Heavy       EVQLVESGGGLVQPGGSLRLSCAASGFTVSSNYMSWVRQAPGKGLEWVSVIYSGGSTF	58
                                         ▪▪▪ ▪▪▪▪                  ▪▪▪▪▪▪▪
                ***********:**************. *****************************:

CV30_Heavy      YADSVKGRFTISRDNSKNTLYLQMNSLRAEDTAVYYCARDLDVSG-GMDVWGQGTTVTVS	117
                                                      ▪▪▪▪▪▪    ▪▪
B38_Heavy       YADSVKGRFTISRHNSKNTLYLQMNSLRAEDTAVYYCAREA---Y-GMDVWGQGTTVTVS	114
                                                      ▪▪▪   ▪   ▪▪
CB6_Heavy       YADSVKGRFTISRDNSMNTLFLQMNSLRAEDTAVYYCARVLPMYGDYLDYWGQGTLVTVS	118
                                                      ▪▪▪▪▪▪▪▪  ▪▪
                *************.** ***:******************        :* ***** ****


CV30_kappa      EIVLTQSPGTLSLSPGERATLSCRASQSVSSSYLAWYQQKPGQAPRLLIYGASSRATG	58
                                           ▪▪▪▪▪▪                       ▪
B38_kappa       DIVMTQSPSFLSASVGDRVTITCRASQGI-SSYLAWYQQKPGKAPKLLIYAASTLQSG	57
                                          ▪▪▪ ▪▪▪
CB6_kappa       DIVMTQSPSSLSASVGDRVTITCRASQSI-SRYLNWYQQKPGKAPKLLIYAASSLQSG	57
                                           ▪  ▪▪▪
                :**:****. ** * *:*.*::*****.: * ** *******:**:****.**:  :*

CV30_kappa      IPDRFSGSGSGTDFTLTISRLEPEDFAVYYCQQYGSS--PQTFGQGTKLEIKRTVAAPSV	116
                          ▪                       ▪▪
B38_kappa       VPSRFSGSGSGTEFTLTISSLQPEDFATYYCQQLNSY-PPYTFGQGTKLEIKRTVAAPSV	116
                         ▪▪▪                    ▪▪▪▪▪   ▪
CB6_kappa       VPSRFSGSGSGTDFTLTISSLQPEDFATYYCQQSYSTPPEYTFGQGTKLEIKRTVAAPSV	117
                         ▪                        ▪▪▪▪▪
                :*.*********:****** *:*****.*****  *     *******************
```
